# The perineurium integrates leptin with its sympathetic outflow to protect against obesity

**DOI:** 10.1101/2024.07.04.602091

**Authors:** Gitalee Sarker, Emma Habermann, Anandhakumar Chandran, Thomas Monfeuga, Cyrielle Maroteau, Sofia Lundh, Andrea Raimondi, Noelia Martínez Sánchez, Bernardo Arús, Matteo Iannacone, Enrique M. Toledo, Ana Domingos

**Author notes:** Correspondence to Ana I. Domingos, Department of Physiology, Anatomy and Genetics, University of Oxford, Oxford, OX31PT.

## Abstract

The regulatory mechanism of leptin’s afferent action in the brain, constituting a negative feedback loop, is contingent upon the efferent sympathetic innervation of white and brown adipose tissues. Nonetheless, the peripheral regulation governing the relative strengths of the afferent and efferent arms remains ambiguous. Using single-cell RNA sequencing on murine sympathetic ganglia, we identified the unique expression of both the leptin receptor (LepR) and the beta 2 adrenergic receptor (Adrb2) in perineurial cells that form a barrier around sympathetic ganglia and nerve bundles in adipose tissues. We show that LepR^+^ Sympathetic Perineurial Cells (SPCs) are molecularly similar to endothelial cells and that conditional knockout of Adrb2 in LepR^+^ SPCs predisposes mice to obesity without affecting food intake. Notably, we found that hyperleptinemia associated with obesity causes apoptosis in SPCs, leading to a significant erosion of the perineurial barrier and concomitant adipose sympathetic neuropathy. We further show that this deleterious effect can be reversed by sympathomimetic beta 2 adrenergic receptor agonism. These results have relevance to human obesity, as we observed a synergistic effect of highly common polymorphisms of *LEPR* and *ADRB2* on the risk of increased BMI in a large European population. We propose that SPCs are the nexus of leptin action by integrating the afferent and efferent arms of the neuroendocrine loop to influence its setpoint.

## Main Text

Leptin regulates body weight via a neuroendocrine negative feedback loop. The hormone leptin is the afferent signal released from white adipocytes, in proportion to fat reserves, and acts on hypothalamic neurons to suppress food intake and activate proportional descending efferent sympathetic activity that triggers in lipolysis in white adipose tissue^1^ and thermogenesis in brown adipose tissue^2–5^. The efferent arm reduces mass via noradrenergic signalling, which in turn reduces leptin levels, thereby closing the loop. However, the cellular and molecular mechanisms regulating the balance between afferent leptin and efferent sympathetic outflow are unknown. Here we identified the sympathetic perineurium as a key player in the neuroendocrine loop of leptin action, regulating the balance between the afferent and efferent arms to control body weight. Sympathetic perineurial cells (SPCs) uniquely express both the LepR and beta2 adrenoceptor (Adrb2), enabling them to sense the balance of the afferent and efferent arms in the neuroendocrine loop of leptin action. High levels of leptin drive apoptosis of SPCs, which can be prevented by sympathomimetic beta 2 adrenergic receptor agonism. The strength of leptin’s efferent sympathetic output is determined by the apoptosis of SPCs and the integrity of the perineurial barrier, which has been shown to have a neuroprotective role^6^. This novel working model could explain why humans with common polymorphisms of both *LEPR* and *ADRB2* have a higher risk of developing obesity and provide a new mechanistic model for the leptin setpoint, one operating beyond the central regulation of appetite.

### 1. Single-cell RNA-seq of sympathetic ganglia reveals a cluster of endothelial cells highly expressing the leptin receptor

To map the heterogeneity of non-neuronal cell types resident in sympathetic ganglia and their plasticity in response to obesity, we performed single-cell RNA sequencing (scRNA-seq) on isolated superior cervical ganglia (SCG) and stellate ganglia from male C57BL/6 mice fed either normal chow diet or high-fat diet (HFD) for 12 weeks. As expected, the HFD fed mice gained substantially more weight compared with the chow fed mice (**Extended Data Fig. 1a**). For each condition, we pooled SCG or stellate ganglia from lean and obese mice prior to scRNA-seq and the four datasets were integrated for analysis (**Fig. 1a**). After low-quality filtering of the scRNA-seq data, we considered a total of 73,042 single cells. After data integration, we performed principal component (PC) analysis and dimensionality reduction with uniform manifold approximation and projection (UMAP) in the integrated dataset^7^. Using unsupervised clustering, we detected 9 distinct clusters of immune and non-immune cells (**Fig. 1b**). Each cluster contained cells from each ganglia type, feeding condition and sample batch, indicating that the transcriptional identities of these cell clusters are stable irrespective of experimental conditions (**Extended Data Fig. 1b, c**). An immune cell marker (*Cd45*) was expressed in all immune cell populations and absent in non-immune cell populations (**Fig. 1c**). Using expression patterns of cell type specific marker genes, we assigned a single identity to each cluster: endothelial cells (*Cd31^+^*), pericytes (*Des^+^*), schwann cells_ non myelinating (*Cdh2*^+^), schwann cells myelinating (*Mbp*^+^), neurons (*Th*^+^), dendritic cells (*Cd11c^+^*), microglia (*Tmem11^+^*), macrophages (*F480^+^*), and T regulatory cells (*Il17r^+^*; **Fig. 1c**). Overall, 74% of the single cell transcriptomes were assigned to glial cells, 10.6% to endothelial cells, 2.1% pericytes, 0.6% neurons and 12.7% to immune cell types (**Extended Data Fig. 1d**).

**Fig. 1.**
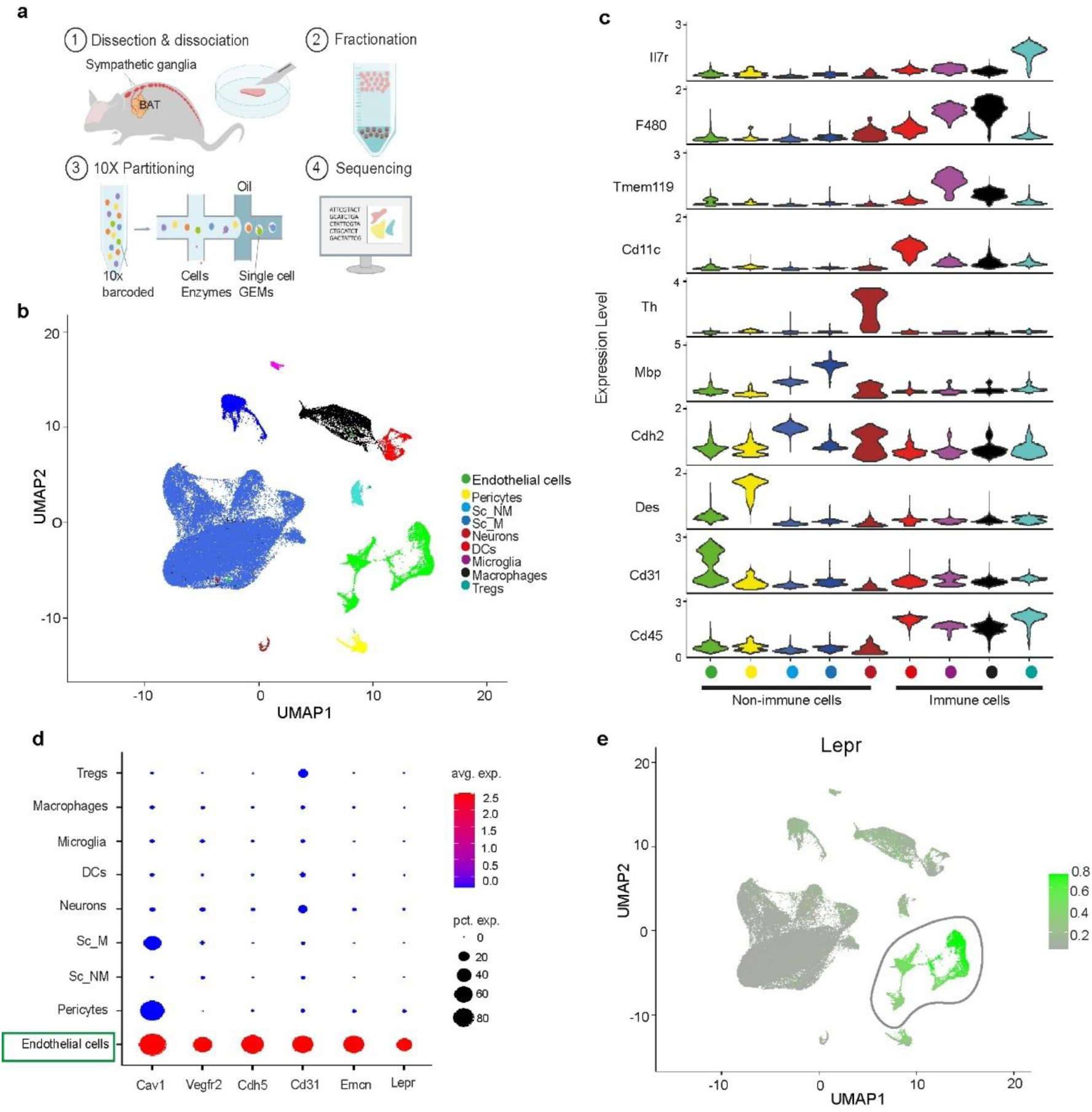
Single-cell transcriptomic data of sympathetic ganglia reveals a unique endothelial population that highly expresses leptin receptor. **a,** Schematic workflow of scRNA-seq of murine sympathetic ganglia. **b,** Uniform manifold approximation and projection (UMAP) plot of all 73,042 cells from mouse superior cervical and stellate ganglia. **c,** Violin plots showing the expression of cell type specific marker genes. The y axis indicates the log normalized gene-expression value and the width indicates the number of cells expressing the particular gene. **d,** Dotplot showing the expression of endothelial markers (Cav1, Vegfr2, Cdh5, Cd31, Emcn) and leptin receptor (Lepr) in endothelial cell population. The size of the dot corresponds to the percentage of cells expressing the gene in each cluster and the color represents the average gene expression level. **e,** Feature plot showing the expression of Lepr in the endothelial cluster. For visualization, denoised expression data was used (c, e). Sc_NM = schwan cell_non myelinating, Sc_M = schwan cell_myelinating, DCs = dendritic cells, Tregs = T regulatory cells.

The endothelial cell cluster exhibited elevated expression of Cav1, Vegfr2, Cdh5, Emcn, Cldn5, Egfl7 that were previously reported to be highly expressed in vascular endothelial cells^8–11^ (**Fig. 1d**). In contrast, we observed very low expression of lymphatic endothelial cell markers (Lyve1, Prox1) in the endothelial cell cluster^10,12^ (**Extended Data Fig. 2a**). Notably, we detected especially high expression of leptin receptor (Lepr) gene in the endothelial cell population (average log2FC = 1.96, percentage of endothelial cells expressing Lepr = 71%, percentage of all other cells expressing Lepr = 0.3%, p < 2.2e-16; **Fig. 1d, e)**. Quantification of Lepr^+^ endothelial cell population revealed that 50.6% of endothelial cells are Lepr^+^ Cav1^+^ (**Extended Data Fig. 2b, c**), 37.8% of endothelial cells are Lepr^+^ Vegfr2^+^ (**Extended Data Fig. 2d, e**) and 47.2% of endothelial cells are Lepr^+^ Cdh5^+^ (**Extended Data Fig. 2f, g**). To visualise the sympathetic endothelial cells that expressed the leptin receptor, we performed immunofluorescence of SCG and sympathetic nerve bundles dissected from the subcutaneous white adipose tissue (scWAT) and brown adipose tissue (BAT) of LepR^Cre^; Rosa26^-Lox-Stop-Lox-ChR2YFP^ reporter (LepR^Cre^;LSL-YFP) mice. In LepR^Cre^;LSL-YFP mice, cre expression follows an IRES sequence downstream of the long form of the leptin receptor^13^ and the cre-mediated recombination results in expression of membrane-bound YFP. Double staining of CAV1, VEGFR2 or CDH5 with YFP in the sympathetic ganglia and sympathetic nerve bundles revealed that YFP is co-expressed with CAV1 (**Extended Data Fig. 3a)**, VEGFR2 (**Extended Data Fig. 3b)** and CDH5 (**Extended Data Fig. 3c)** positive endothelial cells forming a thin layer surrounding the TH^+^ (tyrosine hydroxylase) sympathetic neurons.

### 2. Lepr^+^ sympathetic endothelial cells constitute the perineurial barrier

The unsheathing structure formed by the Lepr^+^ endothelial cells resemble the perineurial barrier, which was once described as epithelial^14^. Interestingly, the scRNA-seq data set reveal that the Lepr^+^ endothelial cell population also highly transcribes Glut1, Itgb4, Lypd2, Mpzl2, Cldn1(**Fig. 2a**) which have been reported as perineurial marker genes in sciatic nerve single cell atlas^10,15^. Among the perineurial markers, Glut1 which has been reported as a pan perineurial marker^16^, was co-expressed predominantly in the Lepr^+^ endothelial cell cluster (**Fig. 2b**). Quantification of Lepr^+^ Glut1^+^ cells among different clusters revealed around 20.2% endothelial cells are Lepr^+^ Glut1^+^ (**Fig. 2c**). The co-expression was also detected *in situ* by GLUT1 and TH staining of SCG and sympathetic nerve bundles innervating the scWAT and BAT of the LepR^Cre^; LSL-YFP reporter mice (**Fig. 2d**). We thus confirmed that the Lepr^+^ sympathetic endothelial cells constitute the perineurial barrier of the sympathetic ganglia and nerve bundles. We, therefore, refer these cells as <u>S</u>ympathetic <u>P</u>erineurial Cells (SPCs).

**Fig. 2.**
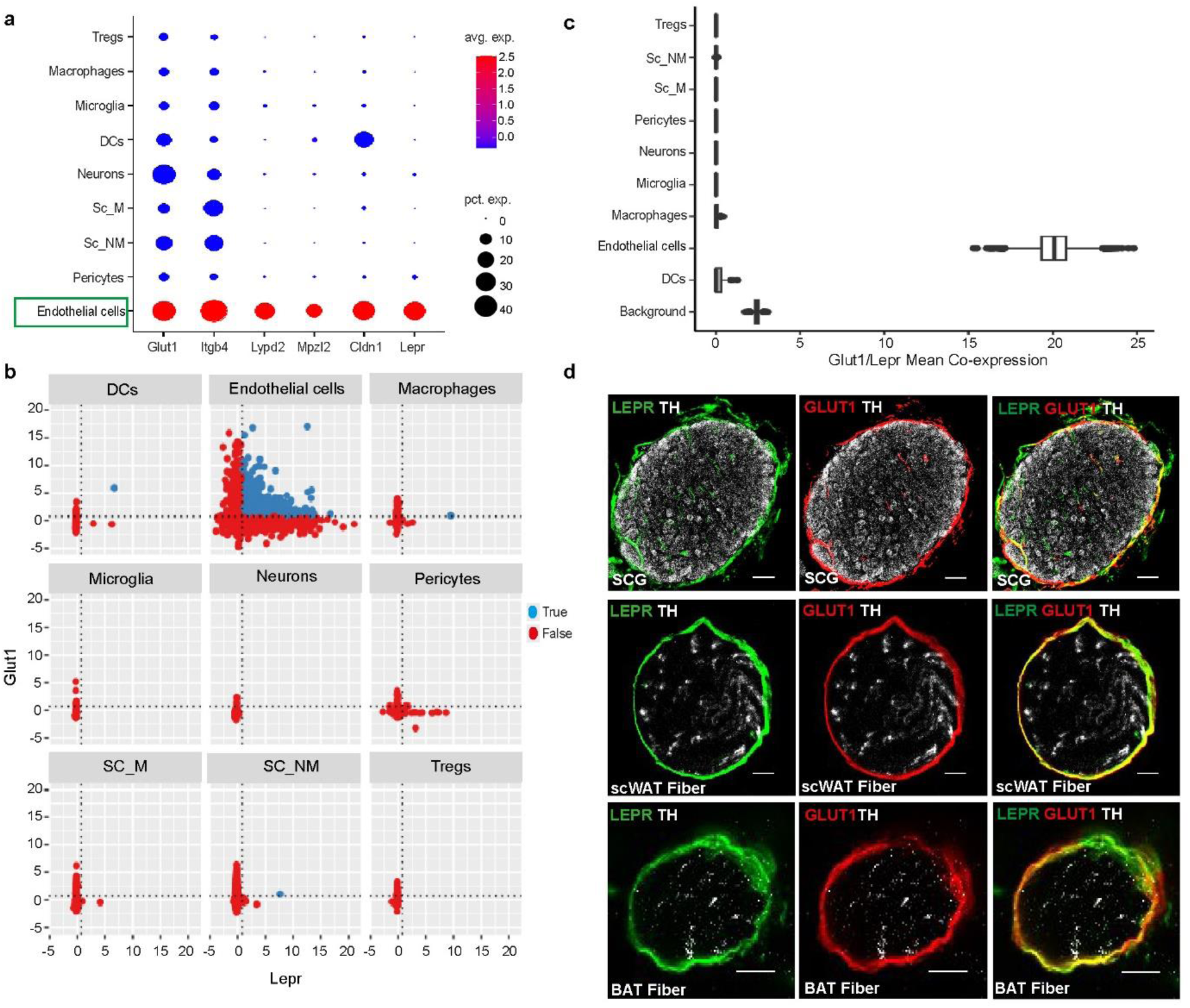
Lepr+ cells surround the sympathetic ganglia and sympathetic nerve bundles in adipose tissue as perineurial barrier. **a,** Dotplot showing the expression of perineurial markers (Glut1, Itgb4, Lypd2, Mpzl2, Cldn1) and leptin receptor (Lepr) in the endothelial cell population. The size of the dot corresponds to the percentage of cells expressing the gene in each cluster and the color represents the average gene expression level. **b,** scRNA analysis shows the co-expression of Lepr and Glut1 in different cell populations. The position of the dotted lines is described in material and methods. **c,** Box plot shows the percentage of cells co expressing Lepr and Glut1. **d,** Immunofluorescence staining of LEPR (YFP), TH and GLUT1 protein in superior cervical ganglia; scale bar = 50 µm, sympathetic nerve bundles innervating the scWAT and sympathetic nerve bundles innervating BAT isolated from 12-week-old male LepR-cre; LSL-YFP reporter mice; scale bar = 20 µm. Sc_NM = schwan cell_non myelinated, Sc_M = schwan cell_myelinated, DCs = dendritic cells, Tregs = T regulatory cells.

### 3. SPCs highly co-express Lepr and Adrb2, unlike any other cell type

We next questioned whether SPCs could sense the sympathetic efferent arm in the neuroendocrine loop of leptin. To answer this question, we first checked in the scRNA-seq dataset whether SPCs express any adrenergic receptors^17^. Among different adrenergic receptors, Adrb2 is highly expressed in the endothelial cell cluster. In contrast, we found very low expression of alpha-adrenergic receptors and no expression of beta 3 adrenergic receptor in the endothelial population (**Extended Data Fig. 4a**). Co-expression analysis further shows that Lepr and Adrb2 are predominantly co-expressed in 30% of the endothelial cell population (**Fig. 3a, b**). To validate the scRNA-seq results, we performed fluorescent *in situ* hybridization in combination with GLUT1 or CAV1 immunohistochemistry by RNAscope to detect the Lepr and Adrb2 mRNA in the SPCs barrier. We observed co-expression of Lepr and Adrb2 in the GLUT1^+^ (**Fig. 3c)** and CAV1^+^ (**Extended Data Fig. 4b)** positive SPCs barrier in SCG and sympathetic nerve bundles.

**Fig. 3.**
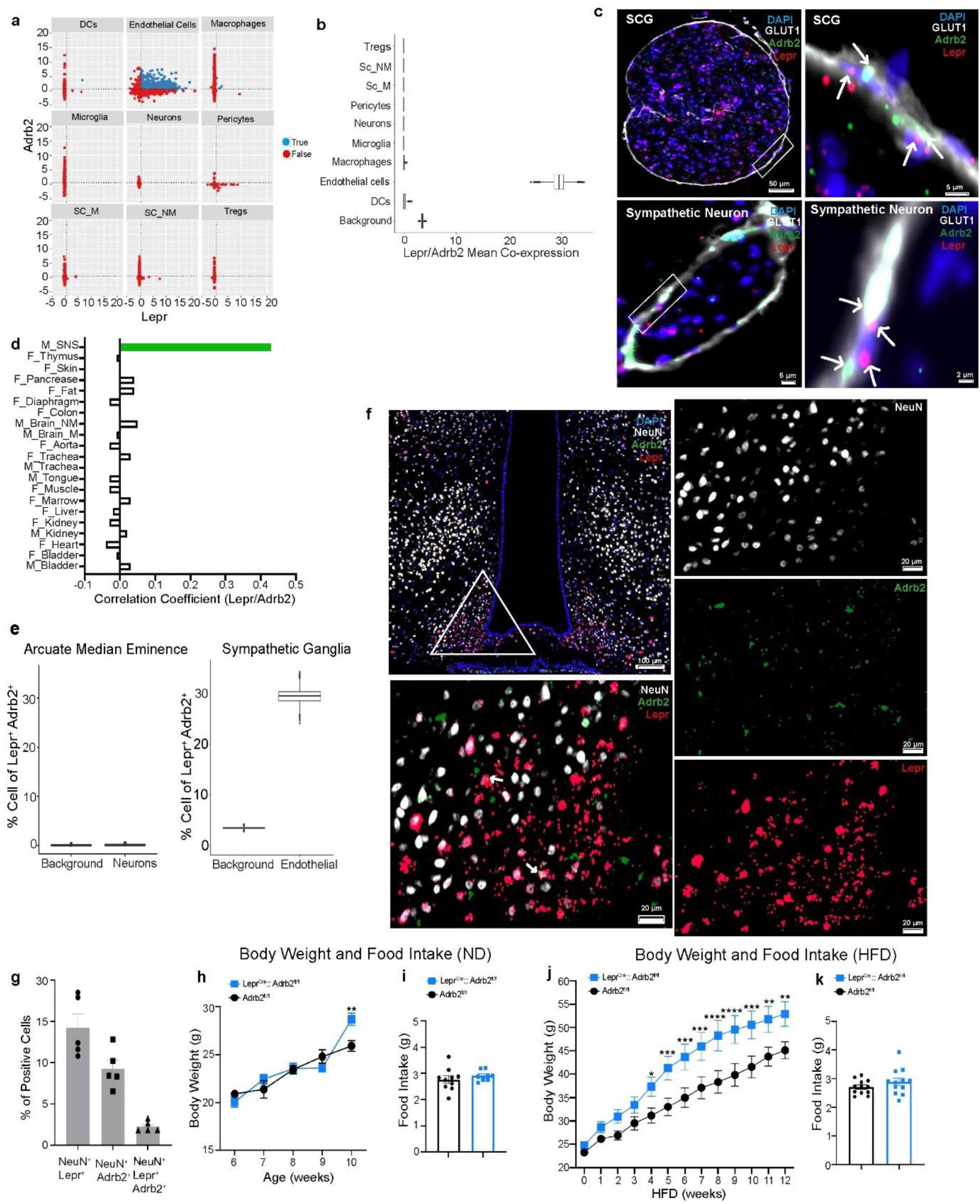
Adrb2 in SPCs is required for metabolic homeostasis. **a,** scRNA analysis shows the co-expression of Lepr and Adrb2 in different cell populations. The position of the dotted lines is described in material and methods. **b,** Box plot shows the percentage of cells co-expressing Lepr and Adrb2. **c,** High magnification image showing the dual labelling of GLUT1 protein with IHC and Lepr, Adrb2 mRNA with ISH in superior cervical ganglia (SCG) and sympathetic nerve bundles dissected from WT adult mice. **d,** Bar plot showing the correlation coefficient of Lepr and Adrb2 in different organs (Tabula Muris data set), M = Microfluidic droplet method and F = FACS based method. **e,** Box plot showing the percentage of Lepr^+^Adrb2^+^ neuronal population in arcuate-median eminence (HypoMap dataset) and endothelial cell population in sympathetic ganglia. **f,** High magnification image showing the dual labelling of NeuN protein with IHC and Lepr, Adrb2 mRNA with ISH in the hypothalamus of WT adult mice. **g,** Bar plot showing the percentage of neurons in arcuate region of hypothalamus expressing Lepr and Adrb2. (n = 5 mice). **h,** Body weight of normal diet (ND) fed Lepr^Cre^; Adrb2^fl/fl^ and control (Adrb2^fl/fl^) mice (n = 17-22 per group). **i,** Food intake of normal diet fed Lepr^Cre^; Adrb2^fl/fl^ and control (Adrb2^fl/fl^) mice (n =17-22 per group). **j,** Body weight of Lepr^Cre^; Adrb2^fl/fl^ and control (Adrb2^fl/fl^) male mice challenged with 12 weeks high fat diet (HFD) (n = 5-6 per group). **k,** Food intake of Lepr^Cre^; Adrb2^fl/fl^ and control (Adrb2^fl/fl^) male mice following 12 weeks high fat diet challenge (n = 5-6 per group). Sc_NM = schwan cell_non myelinated, Sc_M = schwan cell_myelinated, DCs = dendritic cells, Tregs = T regulatory cells. Data were mean ± s.e.m and were analyzed using two-way ANOVA with Bonferroni post-hoc test (h, j) and two tailed unpaired Student’s t-test (i, k). *p < 0.05, **p < 0.01, ***p < 0.001, ****p < 0.0001.

Next, we investigated whether Lepr and Adrb2 were co-expressed in any other tissues. To this end, we estimated Lepr-Adrb2 gene-gene correlation coefficient (r) in the single cell transcriptomic data from 16 different murine organs published in Tabula Muris compendium^18^. We observed weak correlation ((0.02 ≤ *r* ≤ 0.05); bladder, kidney, marrow, trachea, brain, fat and pancreas and (−0.04 ≤ *r* ≤ −0.01); heart, liver, muscle, tongue, aorta, thymus, diaphragm) or no correlation (skin, colon) of this gene pair in the datasets (**Fig. 3d**). In contrast, we found relatively higher correlation (r = 0.43) of these two genes in our scRNA dataset of sympathetic ganglia (**Fig. 3d).** We further analysed the co-expression of this gene pair in the different cell clusters extracted from murine heart, liver, aorta, fat, pancreas, and muscle^18^. Our analysis did not find significant co-expression of Lepr and Adrb2 any other cell type in other organs thus far (**Extended Data Fig. 4c-h**).

Neurons in arcuate nucleus of hypothalamus express leptin and adrenergic receptors which play major role in regulating food intake as well as sympathetic outflow to white and brown adipose tissues^19–23^. We, therefore, analysed the HypoMap scRNA dataset to investigate the possible presence of Lepr^+^ Adrb2^+^ co-expression in the arcuate neuronal populations, which have roles in the regulation of food intake and SNS activity^24^. Our analysis revealed a negligible percentage of neurons (0.01%) in arcuate hypothalamus co-express Lepr and Adrb2 mRNA compared to that of sympathetic endothelial cells (30%) (**Fig. 3e, Extended Data Fig. 5a-c)**. This observation was further validated by dual labelling of Lepr and Adrb2 mRNA in NeuN positive neurons in arcuate hypothalamus by fluorescent *in situ* hybridization and immunohistochemistry (**Fig. 3f)**. The quantification of Lepr^+^ Adrb2^+^ neurons in arcuate hypothalamic region revealed an average of only 2.3 ± 0.3% CNS neurons as Lepr^+^ Adrb2^+^ (**Fig. 3g)**. We also analysed the arcuate neuronal populations within the HypoMap dataset ascertain whether Lepr is co-expressed with other alpha- and beta-adrenergic receptors. Our co-expression analysis excluded the presence of any significant co-expression of Lepr with any other adrenergic receptor subtypes in arcuate neurons **(Extended Data Fig. 5d-k)**.

### 4. Adrb2 in SPCs protect against obesity, independently of food intake

To test whether Adrb2 in SPCs was functionally relevant to body weight homeostasis, we generated Lepr^cre^; Adrb2^fl/fl^ conditional knockout mice where Adrb2 is specifically deleted from Lepr^+^ cells. Although there was no difference in body weight between Lepr^cre^; Adrb2^fl/fl^ and Adrb2^fl/fl^ control mice until early adulthood, by 10 weeks of age Lepr^cre^; Adrb2^fl/fl^ mice kept on normal chow diet weighed more (13% heavier than Adrb2^fl/fl^ control mice) independently of food intake (**Fig. 3h, i)**. Two groups were then challenged with 12 weeks ad libitum HFD from 8 weeks of age. We observed that, Adrb2^fl/fl^ control mice, the body weights of Lepr^cre^; Adrb2^fl/fl^ mice was increased by nearly 30% after 8 weeks on HFD. On average, between weeks 5-12 Lepr^cre^; Adrb2^fl/fl^ mice were 22.3% heavier compared to Adrb2^fl/fl^ control mice (**Fig. 3j)**. The weight gain was not due to excess food intake since they did not display any difference in food intake (**Fig. 3k)**. Thus, the expression of Adrb2 in Lepr^+^ SPCs is necessary to maintain a lower body weight and protect against obesity.

### 5. Diet induced obesity destroys the SPCs barrier

We next asked whether obesity changes the integrity of the SPCs barrier. Analysis of the frequency of cells in the endothelial cluster of lean and obese sympathetic ganglia revealed a decreased proportion of endothelial cells in obese mice (**Fig. 4a**). Comparison analysis of Lepr^+^ endothelial subset between lean and obese states further depicted a 20% reduction of Lepr^+^ endothelial cells in the sympathetic ganglia of obese mice (**Fig. 4b**). To corroborate the scRNA-seq analysis, we immunolabelled the scWAT and BAT sympathetic nerve bundles of 12 weeks DIO and normal chow-fed lean LepR^Cre^; LSL-YFP reporter mice. These studies revealed a significant reduction in Lepr^+^ SPCs area in the sympathetic nerve bundles innervating both the scWAT (**Fig. 4c, e**) and the BAT (**Fig. 4d, g**) of DIO mice compared to aged-matched lean mice. The density of TH^+^ sympathetic axons was also significantly decreased in both scWAT (**Fig. 4c, f**) and BAT (**Fig. 4d, h**) sympathetic nerve bundles of DIO mice. By using electron microscopy on scWAT derived sympathetic nerve bundles of DIO and chow-fed lean mice, we confirmed a significant loss of perineurial cell layers in obese mice compared to their lean littermates (**Fig. 4i, j**). Thus, diet induced obesity disrupted the SPCs barrier, and which is concomitant with sympathetic neuropathy^25,26^ in adipose tissues.

**Fig. 4.**
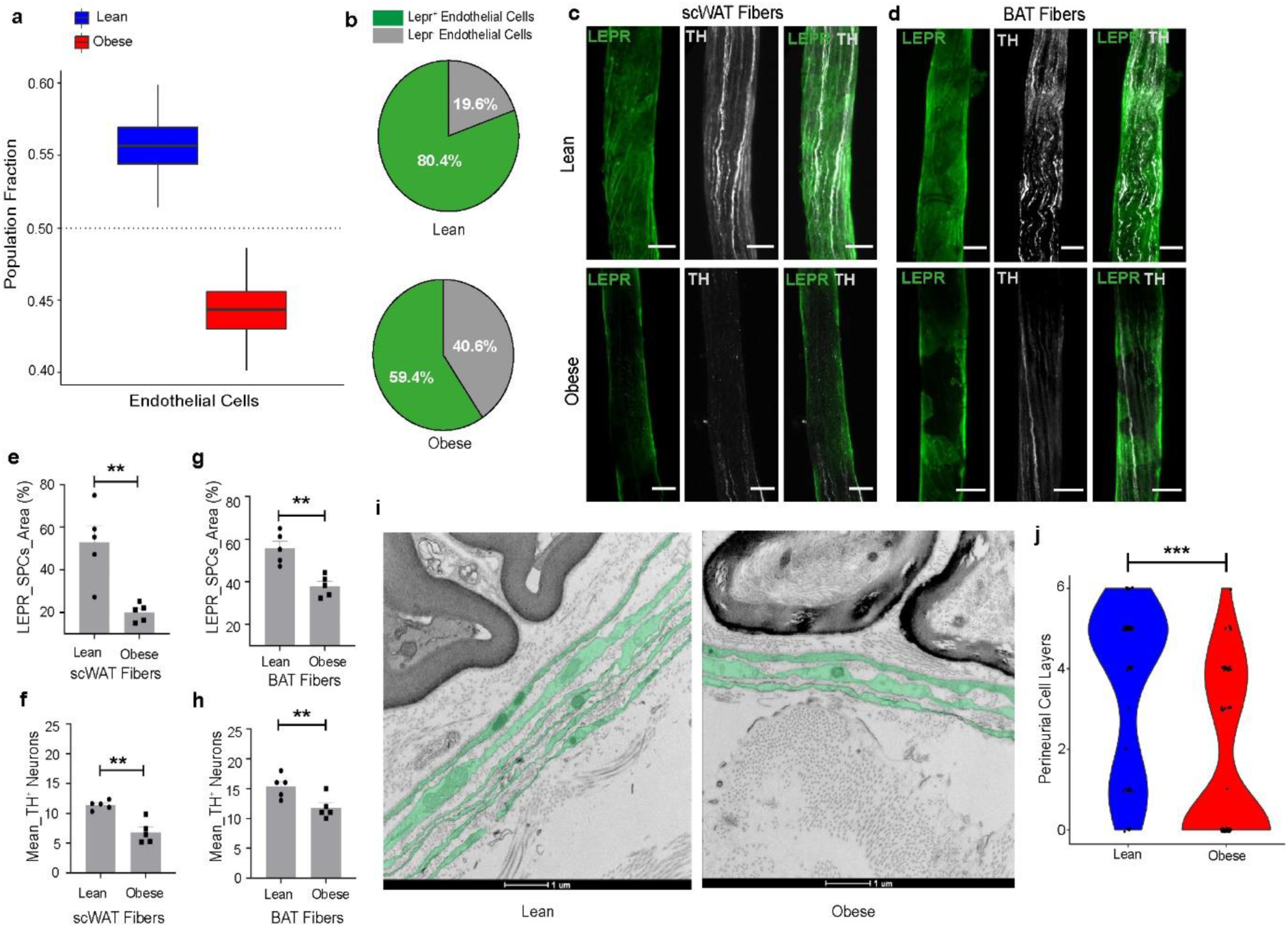
Obesity reduces the SPCs and destroys the perineurial barrier. **a,** scRNA-seq of ganglia showing the fraction of endothelial cell population in normal chow diet (ND)-fed lean and high fat diet (HFD) fed-obese WT mice. **b,** Pie chart shows the percentage of Lepr^+^ endothelial cells in lean and obese endothelial population within ganglia. **c,** Representative images of scWAT bundles from ND-treated lean and HFD treated-obese (LepR^cre^; LSL-YFP) reporter mice showing the expression of LEPR (YFP) and TH. **d,** Representative images of BAT bundles from ND-treated lean and HFD-treated obese (LepR^cre^; LSL-YFP) reporter mice showing the expression of LEPR (YFP) and TH. Scale bar: 50µm. **e-f,** Quantification of LEPR^+^ SPCs barrier and TH^+^ sympathetic neurons in ND-treated lean and HFD-treated obese scWAT bundles (n = 5 mice per group). **g-h,** Quantification of LEPR^+^ SPCs barrier and TH^+^ sympathetic neurons in ND-treated lean and HFD-treated obese BAT bundles (n = 5 mice per group). Data were analyzed by two-tailed unpaired Student’s t-test and were shown as mean ± s.e.m (e-h). **i,** Electron micrograph showing the perineurial barrier in scWAT bundles of lean and obese mice with false coloring to highlight the perineurial layers (green). **j,** Quantification of the perineurial layer in obese and lean scWAT bundles (n = 3 mice per group) and were analyzed by two-tailed unpaired Student’s t-test. Scale Bar: 1µm. *p < 0.05, **p < 0.01, ***p<.001.

### 6. Obesity and high leptin, drives apoptosis of SPCs which is prevented by sympathomimetic Adrb2 agonism

We next asked how diet induced obesity causes loss of perineurial barrier. To answer that question, we performed functional enrichment analysis of the differentially expressed genes (DEG) (p.adj < 0.05) in endothelial cell clusters between obese and lean mice based on KEGG (Kyoto Encyclopedia of Genes and Genomes) pathways using ShinyGO 0.77 (with an FDR cut-off of 0.05) ^27^. The top 20 enriched pathways are represented in the dot plot **(Fig. 5a)**. Notably, KEGG pathway analysis revealed the enrichment of apoptotic process in obese endothelial cluster compared to the lean mice **(Fig. 5a)**. Differential gene expression analysis of endothelial cells show that the genes listed in the cellular apoptotic pathways were mostly upregulated in obese mice **(Fig. 5b).** Hence, we next immunolabelled the SCG and scWAT derived sympathetic nerve bundles of 12 weeks DIO obese and lean mice for one of the apoptotic markers, Tnfrsf1a (a.k.a TNFR1). Consistent with the scRNA-seq analysis, we observed a higher expression of TNFR1 in the SPCs barrier and the sympathetic neurons of SCG **(Fig. 5c)** and scWAT sympathetic nerve bundles **(Fig. 5d, e)** of obese mice compared to their lean littermates. Together, this data suggests that diet induced hyperleptinemic obesity drives apoptosis of SPCs barrier and sympathetic neurons, resulting in loss of SPCs barrier and sympathetic neuropathy in adipose tissue. To ascertain this observation, we performed *ex vivo* experiments by treating SCG explants with low leptin (10 ng/ml) and high leptin (100 ng/ ml) that, respectively, emulate in vitro the states of normo- and hyperleptinemia^25,28^. We then evaluated the effect of high leptin on the CAV1^+^ SPCs barrier and TH^+^ sympathetic neurons by immunofluorescence. Compared to low leptin treatment, we observed a significant reduction of CAV1 expression in the SPCs barrier and TH intensity in the sympathetic neurons of SCG explant after stimulation with high dose of leptin **(Fig. 5f-h)**. We next checked whether activation of Adrb2 in SPCs can rescue the high leptin effect on SCG explant. Notably, we observed that co-incubation with high dose of leptin and a sympathomimetic beta2 agonist could restore the CAV1^+^ SPCs barrier and TH^+^ neurons **(Fig. 5f-i)**. Next, we evaluated the expression of apoptotic marker TNFR1 in the SCG explant treated *in vitro* with different doses of leptin. Consistent with the *in situ* data (**Fig. 5c)**, we observed significantly increased expression of TNFR1 in SPCs barrier and sympathetic neurons following high dose of leptin, which was restored following treatment with the beta2 agonist **(Extended Data Fig. 6).** Collectively, the data suggests that obesity-driven hyperleptinemia causes apoptosis of SPCs and sympathetic neurons and that this effect can be prevented by activating beta 2 adrenergic receptors in SPCs.

**Fig. 5.**
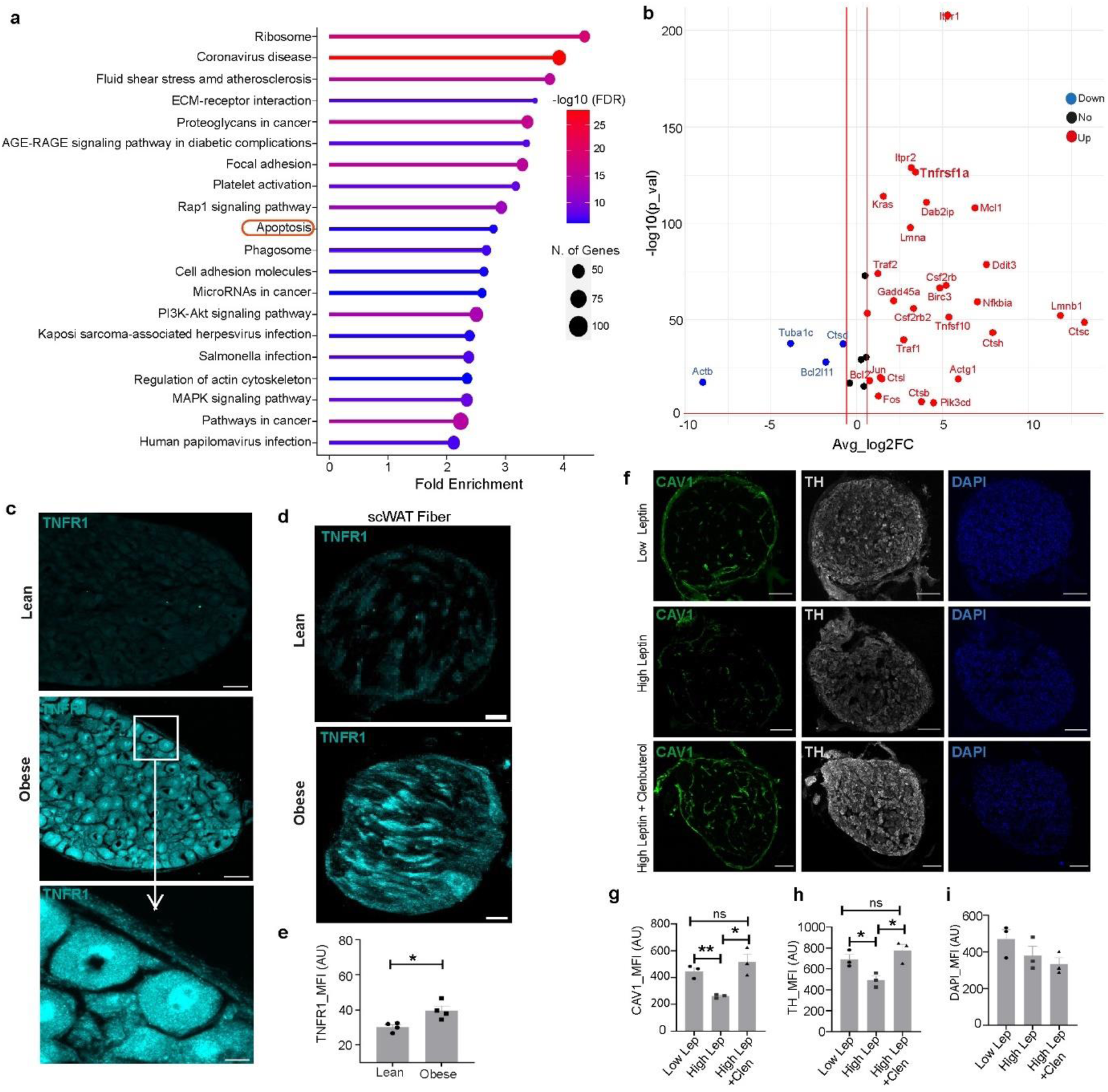
Obesity drives apoptosis of the SPCs which is reversed by beta 2 adrenergic agonism. **a,** The top 20 enriched KEGG pathways using the differentially expressed genes between lean and obese sympathetic endothelial cell population. **b,** Volcano plot showing the differentially expressed genes between lean and obese sympathetic endothelial cell population that are involved in apoptosis. The logarithms of the foldchanges of individual genes (x-axis) are plotted against the negative logarithm of their P-value to base 10 (y-axis). Positive log2 (fold change) values represent up-regulation (red) and negative values represent down-regulation (blue) in obese mice. The red vertical lines represent more than 50% of fold change. **c-d,** Representative images of (c) SCG, Scale bar, 50 µm and (d) scWAT bundles from normal diet (ND)-fed lean and high fat diet (HFD)-fed obese mice showing the expression of TNFR1, Scale bar, 50 µm. **e,** Image quantification of TNFR1 expression in lean and obese scWAT bundles (n = 4 mice per group). **f,** Representative images of SCG explant culture showing the expression of CAV1, TH and DAPI following low leptin (10 ng/ml), high leptin (100 ng/ml) and high leptin (100 ng/ml) plus clenbuterol (10 µg/ml) treatment. Scale bar, 100 µm. **g-i,** Quantification of CAV1, TH and DAPI expression in SCG explant following different stimulating conditions (n = 3 mice per condition). MFI = Mean Fluorescence Intensity. Data are mean ± s.e.m and were analyzed using two-tailed unpaired Student’s t-test (e) and one-way ANOVA with Turkey’s Multiple Comparison test. *p < 0.05, **p < 0.01.

### 7. Common polymorphisms of LEPR and ADRB2 affect BMI synergistically

In humans, recent genetic association studies in middle-aged Brazilian and Japanese population reported a synergic interaction between *LEPR-ADRB2* variants for being associated with the risk of overweight/obesity^29,30^. We analysed such *LEPR* and *ADRB2* variants effect on the European individuals from the UK Biobank cohort. Consistent with these reports, we observed that a *LEPR* variant Gln223Arg (rs1137101) or an *ADRB2* variant Gln27Glu (rs1042714) do not have, individually, any statistically significant effects on BMI (**Table 1**). However, we found a significant synergistic effect between *LEPR* Gln223Arg (rs1137101) variant and *ADRB2* Gln27Glu (rs1042714) variant for BMI, (beta = 0.04; P value = 0.013 (**Table 1**)). The same interaction was with a higher effect in the BMI > 25 group (beta = 0.047, P value = 0.0056) but not significant in the BMI < 25 group (**Table 1**). Together, the genetic data points to an interaction between *LEPR* and *ADRB2* variants in the association with BMI within the European population (interaction frequency ∼33%) and replicate the findings within the Brazilian and Japanese population.

**Table 1.** The effects of *LEPR* and *ADRB2* variants in the regulation of BMI. *LEPR* (rs1137101) and *ADRB2* (rs1042714) effects on BMI. EA = Effect Allele, EAF = Effect Allele Frequency, AA = Amino Acid, Beta = Effect size, SE = Standard Error, IAF = Interaction Allele Frequency, *P* = P value. Data were analysed using a linear regression with an interaction variable corrected for age, sex and 10 first principal components with Bonferroni correction for multiple testing. *P value<0.025.

|  |  |  |  |  |  |  |  |  | Overall European Population |  |  | European Population BMI<25 |  |  | European Population BMI >25 |  |  |
| --- | --- | --- | --- | --- | --- | --- | --- | --- | --- | --- | --- | --- | --- | --- | --- | --- | --- |
| Gene | Variant | EA | EAF | AA change | Consequence | Beta | SE | P | IAF | Interaction effect in Europe unrelated |  | IAF | Interaction effect in Europe unrelated |  | IAF | Interaction effect in Europe unrelated |  |
|  |  |  |  |  |  |  |  |  |  | Beta | P |  | Beta | P |  | Beta | P |
| *LEPR* | rs1137101 | G | 0.5 | Gln223Arg | Missense | -0.99 | 2.72 | 0.71 | 0.33 | 0.0407 | 0.013* | 0.30 | 0.0408 | 0.037 | 0.36 | 0.047 | 0.0056* |
| *ADRB2* | rs1042714 | C | 0.68 | Gln27Glu | Missense | -0.0051 | 0.011 | 0.66 |  |  |  |  |  |  |  |  |  |

## Discussion

Systems control theory define that the setpoint of negative feedback loop is attained when the magnitudes of the afferent and efferent arms act in proportion. To this end, the loop ought to have an integrator, that senses both the afferent and efferent arms, and that determines the operating balance. In searching for a cell that could sense both afferent and efferent arms in the neuroendocrine loop of leptin, we encountered in sympathetic ganglia and nerve bundles in adipose tissues, a previously undescribed perineurial endothelial cell population (SPCs) that uniquely expresses both the Lepr and the Adrb2. Such co-expression was not observed in other cell type in any other organs analysed so far^18^. Thus, Lepr^+^ Adrb2^+^ SPCs are well poised to sense and integrate leptin and NE in adipose tissues^3^. In obesity, an imbalance between the efferent and afferent axis progressively emerges upward shifting the setpoint. To date, the only plausible explanation for this phenomenon was central leptin resistance. We put forward an alternative, if not complementary, explanation that bypasses the brain by acting directly on the efferent sympathetic arm: SPCs work as a peripheral rheostat that integrates both leptin and its descending sympathetic output. When leptin signalling exceeds the one that is triggered by noradrenaline released by sympathetic nerves, SPCs undergo apoptosis which erodes the neuroprotective perineurial barrier, exposing sympathetic neurons to pro-inflammatory cues^31^. Consistent with our observations, a recent study reported how chronic hyperleptinemia worsens diet induced obesity^32^. They showed that a partial reduction of leptin levels (either by genetic strategies or by leptin neutralizing antibodies) can restore leptin sensitivity and facilitate weight loss^32^. Notably, we show that loss of SPCs and degeneration of sympathetic neurons driven by supraphysiologic levels of leptin is completely prevented by beta 2 adrenergic agonism, which mimics sympathetic outflow. This rescue place SPC at a nexus of afferent and efferent arms in the neuroendocrine loop of leptin action.

Consistent with this notion, conditional knockout of Adrb2 in Lepr^+^ SPCs quickly precipitates obesity in mice metabolically challenged with a HFD and promotes adult-onset obesity even when mice ate a regular chow diet. Notably, high fat diet accelerates the weight gain in these mice without affecting their food intake. This metabolic phenotype emphasizes the essential role of Adrb2 in SPCs in regulating body weight. In particular, these results provide new insights onto Adrb2’s regulation of metabolic rate and fat mass reduction, which so far focus on postsynaptic effects^33^ on thermogenic adipocytes, whereas SPCs act presynaptically by unsheathing sympathetic ganglia and nerve bundles, to preserve their normal function^6^.

We further assessed the significance of *LEPR-ADRB2* axis in human obesity, by probing genetic associations in a large European population. We replicated a synergistic interaction of *LEPR* and *ADRB2* variants, on the risk of obesity that was also observed in smaller Japanese and Brazilian cohorts^29,30^. Such synergistic interaction between *LEPR* and *ADRB2* polymorphisms on BMI could likely be explained by altered receptor sensitivity or downstream integration of signalling pathways^34^. Future work will determine how LEPR and ADRB2 signalling cell-autonomously interact to trigger SPCs apoptosis. This will provide new insights on how to dually target LEPR and ADRB2, potentially with unimolecular polypharmacy, to bypass central leptin resistance beyond suppression of food intake in a large patient cohort.

## Methods

### 1. Antibodies, reagents, and drugs

Antibodies were obtained from the following vendors: rabbit anti-tyrosine hydroxylase (TH) (Millipore, Cat #AB152), chicken anti-TH (Aves Lab, Cat #TYH, lot TH1205), goat anti-GFP (Abcam, Cat # ab6673), chicken anti-GFP (Abcam, Cat # ab13970), rabbit anti-caveolin-1 (CAV1) (Cell Signaling, Cat #D46G3), rabbit anti-glucose transporter (GLUT1) (Abcam, Cat #ab150299), rabbit anti-TNFR1 (Invitrogen, Cat #PA595585), rabbit anti-VE-cadherin (Life Technologies Ltd, Cat #361900), goat F (ab) anti-mouse IgG (H+L) (Abcam, Cat #ab6668), mouse anti-VEGFR2 (Santa Cruz Biotechnology, Cat #Sc-6251), rabbit anti-NeuN (Abcam, Cat #ab177487), goat anti-rabbit IgG (H+L) secondary antibody, Alexa Fluor 647 (Invitrogen, Cat #A11010), goat anti-rabbit IgG (H+L) secondary antibody, Alexa Fluor 546 (Invitrogen, Cat #A11035), goat anti-chicken IgG (H+L) secondary antibody, Alexa Fluor 594 (Invitrogen, Cat #A11042), goat anti-chicken IgY (H+L) secondary antibody, Alexa Fluor 647 (Invitrogen, Cat #A21449), goat anti-chicken IgY (H+L) secondary antibody, Alexa Fluor 546 (Invitrogen, Cat #A11040), goat anti-mouse IgG (H+L) secondary antibody, Alexa Fluor 594 (Invitrogen, Cat #A11005), goat anti–chicken IgY (H+L) secondary antibody, Alexa Fluor 488 (Invitrogen, Cat #A-11039), goat anti–rabbit IgG (H+L), Alexa Fluor 488, (Invitrogen, Cat #A11034), donkey anti-goat IgG (H+L) secondary antibody, Alexa Fluor 647 (Invitrogen, Cat #A21447), donkey anti-chicken IgY (H+L) secondary antibody, Alexa Fluor 647 (Stratech, Cat #703-605-155), donkey anti-goat IgG (H+L) secondary antibody, Alexa Fluor 488 (Invitrogen, Cat #A11055), donkey anti-chicken IgG (H+L) secondary antibody, Alexa Fluor 488 (Jackson ImmunoResearch, Cat #703-545-155), donkey anti-rabbit IgG (H+L) secondary antibody, Alexa Fluor 546 (Invitrogen, Cat #A10040), DAPI (Invitrogen, Cat #D1306). Recombinant mouse leptin was obtained from Amylin Pharmaceuticals (R&D Systems, Cat #498-OB). Clenbuterol hydrochloride was purchased from Sigma-Aldrich (Cat #C5423).

### 2. Animals

C57BL/6 wild-type mice at 6-8 weeks old were purchased from Charles River. LepR-cre mice (Lepr^tm2(cre)Rck^; stock no. 008320), Rosa26-LSL-ChR2-YFP mice (stock no. 012-569) were purchased from the Jackson Laboratory and Adrb2^flox/flox^ mice was kindly provided by Gerad Karsenty, Columbia University, USA. Mice were bred in-house to produce homozygous LepR-cre; Rosa26-LSL-ChR2-YFP mice and LepR-cre; Adrb2^flox/flox^ mice. Mice were housed in groups of 2-5, unless fighting or barbering was observed. The room in which animals were housed had a 12-hour light/dark cycle: light during the day and dark during the night, switching at 7:00 and 19:00. Ambient room temperature was maintained at 22 ± 2 °C and humidity at 50 ± 20%. Mice had unrestricted access to water and a regular chow diet (63% carbohydrate, 23% protein and 4% fat), unless mentioned otherwise. Male mice were used in this study. All experiments were conducted in accordance with the United Kingdom Animal Scientific Procedures Act 1986 under personal and project licences granted by the United Kingdom Home Office and approved by the local Department of Physiology Anatomy and Genetics (University of Oxford) ethical review committee.

### 3. High fat diet challenge

When C57BL/6 wild type mice or both LepR-cre; Rosa26-LSL-ChR2-YFP mice, LepR-cre; Adrb2^flox/flox^ mice and respective controls LepR-cre and Adrb2^flox/flox^, reached 8 weeks of age, regular chow diet was replaced with high fat diet (HFD) (20% carbohydrate, 20% protein, 60% fat; Research Diets, Inc., D12492). Length of exposure to HFD is mentioned in figure legends. The body weight and food intake were measured weekly from the onset of high fat diet challenge.

### 4. Tissue harvest and dissociation

Single cell RNA sequencing was performed as previously described^35^.10 adult C57BL/6 males were sacrificed at 24 weeks of age and sympathetic ganglia (SCG and Stellate) were quickly extracted under a stereomicroscope. Tissue was digested for 30 min with collagenase (2.5 mg/ml) and DNase (5 U/ml) in HBSS at 37 ◦C, washed and further digested with trypsin (0.25%) for 30 min at 37◦C with shaking. The samples were mechanically triturated, and the cell suspension was then filtered through a 40-μm cell strainer and centrifuged at 400 g for 5 min. After aspiration of the supernatant, the pellet was resuspended in DMEM media with 10% FBS and 1 vol. of 40% w/v iodixanol solution. The cell suspension was carefully overlayed with an Optiprep gradient composed of 3 ml 22% w/v iodixanol solution and 0.5 ml DMEM and centrifuged at 800 g for 25 min. The viable cells were carefully collected from the top interface and the cell suspension was concentrated to 300 µl by removing the supernatant.

### 5. Library preparation, sequencing and alignment

The cell suspension (approximately 10,000 cells per channel) was then loaded onto the single cell 3’ chip and placed on a 10X Genomics chromium controller instrument to generate single cell gel beads in emulsion (GEMs). Single cell RNA-seq libraries were prepared using the chromium single cell 3’ library & cell bead kit according to the manufacturer’s protocol. Libraries were sequenced with an Illumina NextSeq500 platform to a depth of approximately 300 million reads per library with 2 × 50 read length. The Cell Ranger Single Cell Software Suite v.2.0.1 was used to perform sample de-multiplexing, alignment, filtering, and UMI counting.

### 6. Cell clustering and cell-type annotation

The cluster identities and filtered gene matrices generated by Cell Ranger software were used as input into the open-source R toolkit Seurat (v.4.1.2) (http://satijalab.org/seurat/)^36^ to produce UMAP, feature plots, violin plots of the mean and variance of the mean and variance of gene expression density. Cell barcodes with <500 transcripts detected or >25% mitochondrial gene expression was first filtered out as low-quality cells. The gene counts for each cell were divided by the total gene counts for the cell and multiplied by a scale factor of 10,000, then log2 transformation was applied to the counts. The FindVariableFeatures function was used to select variable genes with default parameters. The ScaleData function was used to scale and center the counts in the dataset. Principal component analysis was performed on the variable genes, and 20 principal components were used for cell clustering (resolution = 0.5) and UMAP dimensional reduction. The cluster markers were found using the FindAllMarkers function, and cell types were manually annotated based on the cluster markers in literature. Module scores were calculated using the AddModuleScore function with default parameters and used to validate certain cell-type annotations. To calculate the sample composition based on cell type, the number of cells for each cell type from each sample were counted. The counts were then divided by the total number of cells for each sample and scaled to 100% for each cell type. The co-expression analysis, the cells having the expression threshold for both genes more than 0.75 are considered to be positive (True) for normalized co-expression. For visualization of gene expression, count data for each run was denoised using a deep count autoencoder, DCA^37^ with default parameters; to be then merged before plotting.

### 7. Immunofluorescence microscopy

Mice were perfused with 1x PBS, sympathetic ganglia and nerve bundles were extracted under the stereomicroscope and fixed in 4% paraformaldehyde overnight at 4°C with shaking. For images in Figure 1d, 4c, d and f, we used frozen sections, and the fixation step was followed by cryoprotection in 30% sucrose (Alfa Aesar). 15-µm sections were obtained in a Leica Cryostat CM3050 S. For images in Figure 3c and 3e, whole-mount tissues were used. Both frozen sections and whole-mount tissues were washed with PBS for 10 min which was followed by incubation in a blocking and permeabilization solution (3% BSA, 2% goat or donkey serum based on the host species of secondary antibodies, 0.1% Tween, and 0.1% sodium azide in 1× PBS) for 1 h at room temperature (RT) with (whole mounts) or without (frozen sections) shaking. The samples were then incubated with primary antibodies overnight at 4 °C with (whole mounts) or without (frozen sections) agitation. The following dilutions of primary antibodies were used: anti-GFP (1:1000), anti-TH (1:500), anti-CAV1 (1:1000), anti-GLUT1 (1:200), anti-TNFR1 (1:500), anti-VEGFR2 (1:500), anti-TUBB3 (1:1000), anti-VE-cadherin (1:100). The anti-GFP antibody was used to detect the YFP tagged LEPR cells. After washing, secondary antibodies (1:500) were applied for 1h at RT, with or without (in the case of frozen sections) shaking. After secondary antibodies incubation, tissues were washed with 1xPBs and was followed by incubation with DAPI (1:1000) for 5 min (frozen sections) or 30 min (whole mounts). After DAPI staining, tissues were washed and mounted with ProLong^TM^ Gold Antifade reagent (Life Technology, Cat #P36930). The slides were dried overnight at RT and stored at 4°C.

Images were acquired with Zeiss LSM 780 Inverted Confocal Microscope. Analysis and quantification of acquired images were performed using FIJI^38^. For representative images, linear adjustments were made to brightness and contrast, whereas for quantification, all channels were kept unchanged. Two to four randomly selected samples per mouse from three to five independent mice per group were quantified.

### 8. Electron microscopy

Sympathetic nerve bundles dissected from subcutaneous white adipose tissue of lean and high fat diet fed obese mice were fixed with 2% paraformaldehyde and 2.5% glutaraldehyde (Electron Microscopy Services) in 0.1 M in Sodium Cacodylate buffer. After several rinses in Sodium Cacodylate buffer samples were postfixed in 1% osmium tetroxide, 1.5% potassium ferricyanide in 0.1 M Sodium Cacodylate buffer for 2 hours on ice. Samples were dehydrated with a crescent concentration of ethanol, washed with propylene oxide and infiltrated in a mixture of propylene oxide/epoxy resin overnight. The resin was then substituted with fresh epoxy resin and samples were embedded in silicone moulds. Then, after being cured for 48 hours at 60°C, resin blocks were cut into ultrathin sections (70 – 90 nm) using an ultramicrotome (UC7, Leica microsystem, Vienna, Austria), longitudinal sections of nerve bundles were collected on copper formvar carbon-coated slot grids, stained with uranyl acetate and Sato’s lead solutions and observed in a Transmission Electron Microscope Talos L120C (FEI, Thermo Fisher Scientific) operating at 120kV. Images were acquired with a Ceta CCD camera (FEI, Thermo Fisher Scientific). For quantification of perineurial layers for each fiber 20 random images were acquired along external fiber profile and quantified manually using ImageJ software^38^.

### 9. Sympathetic ganglia explant cultures

Ganglia explant culture was performed as previously described^31^. SCG and stellate ganglia were removed from mice aged 8 weeks under a stereomicroscope and placed in DMEM (Invitrogen, Carlsbad, CA, USA). Ganglia were cleaned from the surrounding tissue and transferred into eight-well tissue culture chambers (Sarstedt, Nümbrecht, Germany) that were previously coated with poly-d-lysine (Sigma-Aldrich, Steinheim, Germany) in accordance with the manufacturer’s instructions. Ganglia were then covered with 5 µl of Matrigel (BD Bioscience, San Jose, CA, USA) and incubated for 7 min at 37 °C. The culture media prepared with DMEM without phenol red (Invitrogen), 10% FBS (Invitrogen), 2 mM L-glutamine (Biowest, Nuaillé, France), nerve growth factor (1:1000, Sigma-Aldrich), and I% Penicillin-Streptomycin was subsequently added. 12 ganglia explant cultures (6 SCG & 6 stellate) were prepared per condition. Ganglia were cultured for a minimum of 24 h before further manipulation. After 24 h, ganglia cultures were washed once with the culture media. The stimulation protocol in Figure 5 was performed for 24 h with the following concentrations of drugs: low leptin (10 ng/ml), high leptin (100 ng/ml)^25^, high leptin (100 ng/ml) + a sympathomimetic drug (Clenbuterol hydrochloride; 10 µg/ml)^39^. Next day, the cultures were washed once with ice cold 1x PBS for 5 min. The explant cultures were then fixed with 4% paraformaldehyde overnight at 4 °C with agitation, cryoprotected in 30% sucrose and subsequently sectioned by cryostat for immunostaining.

### 10. RNAscope dual *in situ* hybridization (ISH) and immunofluorescence (IF)

Mice at 8 weeks age were perfused transcardially with PBS (pH 7.4). Tissues (brain, SCG, stellate and sympathetic nerve bundles) were immediately removed from the perfused mice and fixed in 10% neutral buffered formalin (Sigma Aldrich) overnight at room temperature (RT) before being transferred to 70% ethanol. The hypothalamic region was isolated from the whole fixed brain as described previously^40^. Samples were dehydrated and embedded in paraffin using standard procedures and then sectioned at 5 µm on to SuperFrost Plus slides (Fisher Scientific). Dual ISH-IF was performed using the RNAscope™ LS Multiplex Fluorescent Reagent Kit (Cat #323275) and RNA-Protein Co-detection Ancillary Kit (Cat #323180) together with Opal 520/570/690 Fluorophore Reagent pack detection (1:1000, Cat #FP1487001KT/FP1488001KT/FP1497001KT) on the Leica BOND RX Fully Automated Research Stainer (Leica, Buffalo Grove, IL) according to the manufacturer’s instructions. The primary antibodies used for IF were rabbit anti-CAV1 (1:1000), rabbit anti-GLUT1 (1:200), rabbit anti-TH (1:1000) and rabbit anti-NeuN (1:500) combined with a Brightvision goat anti-rabbit poly-HRP detection system (Immunologic, NL). For ISH, the following RNAscope target probes from ACD were used: LepR (Mm-Lepr-tv1, ACD Cat #471178) and Adrb2 (Mm-Adrb2-XRn-C2, ACD Cat #1172758-C2), positive control probes: Th (Mm-Th-XRn-C2, ACD Cat #870118-C2) and Polr2a (Mm-Polr2a, ACD Cat #312478), or negative control probe DapB (ACD Cat #312038) Slides were counterstained with spectral DAPI and then cover slipped with Prolong^TM^ Gold Antifade reagent (ThermoFisher Scientific).

Fluorescent slide scans were acquired with an Olympus VS200 slide scanner (Olympus) using a 40x (0.95 NA) air objective and a DAPI/CY3/CY5 filter set. Images were visualized and processed with Olympus OlyVIA 3.8 software. Image quantification was performed using FIJI^38^. Cell counting was performed manually. To quantify the Lepr^+^Adrb2^+^ neurons in arcuate hypothalamus, images were captured from 3 sections per mouse and a total five mice were independently analyzed. The percentage of Lepr^+^, Adrb2^+^ or Lepr^+^Adrb2^+^ positive cells in NeuN^+^ neurons were calculated by total fluorescence positive Lepr, Adrb2 or Lepr Adrb2 mRNA counts, divided by NeuN positive cells, respectively. Percent positive cell values were imported into Prism (GraphPad) for graphing and statistical analysis.

### 11. Quality Control in the UK Biobank (UKB)

*LEPR* and *ADRB2* variants were extracted from the Whole Exome Sequencing (WES) data from 470K UK biobank individuals. Variants were required to pass the following criteria to be selected: 10x minimum coverage, genotype quality score (GQ) ≥ 20, quality score (QUAL) ≥ 30, 0.2 ≤ heterozygote alternative allele ratio ≤ 0.8, 0.8 ≤ homozygote alternative allele ratio ≤1.0, mapping quality score (MQ) ≥ 40 and DPGLnexus variant status = PASS. Furthermore, for both genes, variants with MAF<0.01 and Hardy-Weinberg Equilibrium (HWE) *P* > 1e-12 or variants with MAF >= 0.01 and HWE *P* > 1e-6, outside of the targeted region: CCDS boundaries (https://www.ncbi.nlm.nih.gov/projects/CCDS/CcdsBrowse.cgi) and with a call rate (CR) < 0.9 were removed before analysis.

### 12. Interaction between *LEPR* and *ADRB2* in UKB

To mitigate a possible increase of variance estimates due to relatedness, we sought to remove individuals up to third-degree relatives from our analyses using KING (v2.2.3)^41^. We then selected a total of 346,177 European unrelated individuals with BMI measures to investigate gene-gene interaction analyses between *LEPR* and *ADRB2* genes. Qualifying SNPs were tested for their association with the continuous biomarker using a linear regression with an interaction variable and adjusted for age, sex, array and PC1-10 using SNPassoc R package^42^. The interaction model used was: Trait ∼ β_0_ + β_1_.SNP_1_ + β_2_.SNP_2_ + β_3_.SNP_1_.SNP_2_ + β_n_.covariate_n._ Significance threshold for interaction was defined by traits using a Bonferroni correction p = 0.05/ (n SNP).

### 13. Statistics

Statistical analyses were performed with GraphPad Prism software (GraphPadPrism 9.3.1) using unpaired Student’s t-test (two-tailed) to compare two groups, one-way ANOVA with Turkey’s post-hoc test to compare more than two groups, and two-way ANOVA with Turkey’s post-hoc test to compare two-factor study design. The data are shown as mean ± s.e.m. unless stated otherwise. For all tests, *P* < 0.05 was considered significant. All data points in the manuscript represent individual biological. All ex-vivo and in-situ experiments were performed at least two times. No statistical methods were used to pre-determine sample size, but the sample size was like those generally employed in the field. Randomization was used whenever possible.

## 14. Data availability

The raw single-cell sequencing data of sympathetic ganglia, supporting the findings in this study, will be available upon request. The raw data of the public dataset re-analysed in this study can be found in the GEO repository with the accession codes tabula GSE109774 and GSE93374.

## 15. Code availability

No custom code or mathematical algorithm was created in this work. A repository for reproducible figures will be available in a public repository.

## Acknowledgement

We thank Jeanette Bannebjerg Johansen (Novo Nordisk A/S) for her technical assistance with RNAscope; the Micron Bioimaging Facility for the confocal microscopy, the members of Domingos Lab and NNRCO for discussions and advice on the project. We thank ALEMBIC facility at San Raffaele Scientific Institute, Milan, Italy for support in electron microscopy analysis. Gitalee Sarker is funded by a Novo Nordisk Postdoctoral Fellowship run in partnership with the University of Oxford.

## Contributions

A.I.D. and G.S. conceived the study. G.S., A.K. and T.M. performed single-cell RNA-sequencing. E.M.T and G.S. analyzed scRNA seq data. S.L. and G.S. designed and did the dual IHC and RNAscope. G.S. designed and performed immunohistochemistry, explant culture, confocal microscopy. E.H. did weight acquisition and food intake of LepR^cre^Adrb2^fl/fl^ mice. C.M. performed genetic analyses on UKB data. A.R. and M.I. performed electron microscopy on samples prepared by N.M.S, and B.A. G.S. and A.I.D. wrote the manuscript with input from all authors.

