## supplementary files for "The perineurium integrates leptin with its sympathetic outflow to protect against obesity"

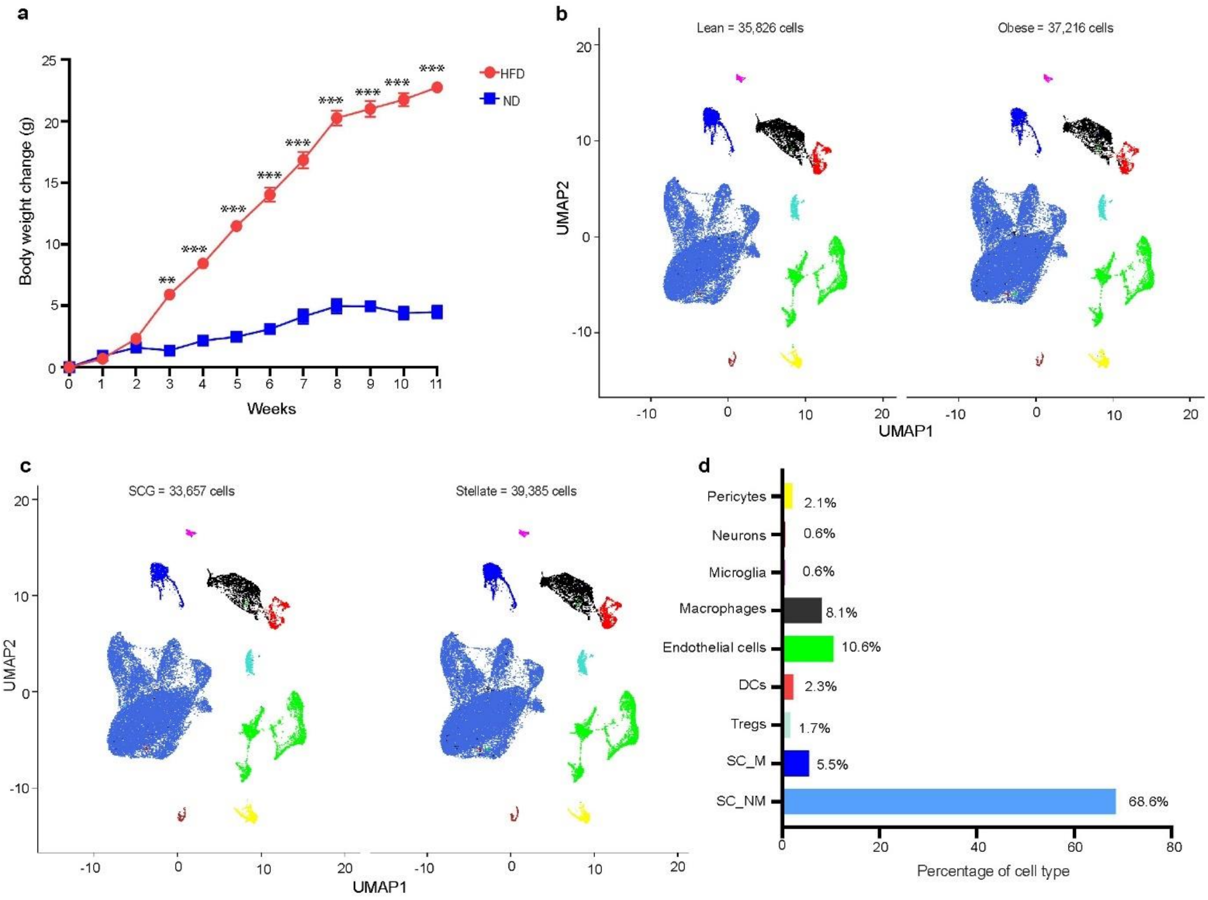


**Extended Data Fig. 1| Single-cell transcriptomic data of sympathetic ganglia from lean and obese mice. a,** The weekly body weight change of WT male mice treated with normal chow diet (ND) and high fat diet (HFD) for 11 weeks (n = 12 per group). **b,** Uniform manifold approximation and projection (UMAP) plot indicating sample classification as lean and obese. **c,** Uniform manifold approximation and projection (UMAP) plot indicating sample classification as superior cervical ganglia (SCG) and stellate. **d,** Bar graph showing the percentage of different cell population. Sc_NM = schwan cell_non myelinating, Sc_M = schwan cell_myelinating, DCs = dendritic cells, Tregs = T regulatory cells. Data were mean ± s.e.m and were analyzed using two-way ANOVA with Turkey’s post-hoc test. **p < 0.01, ***p<0.001.

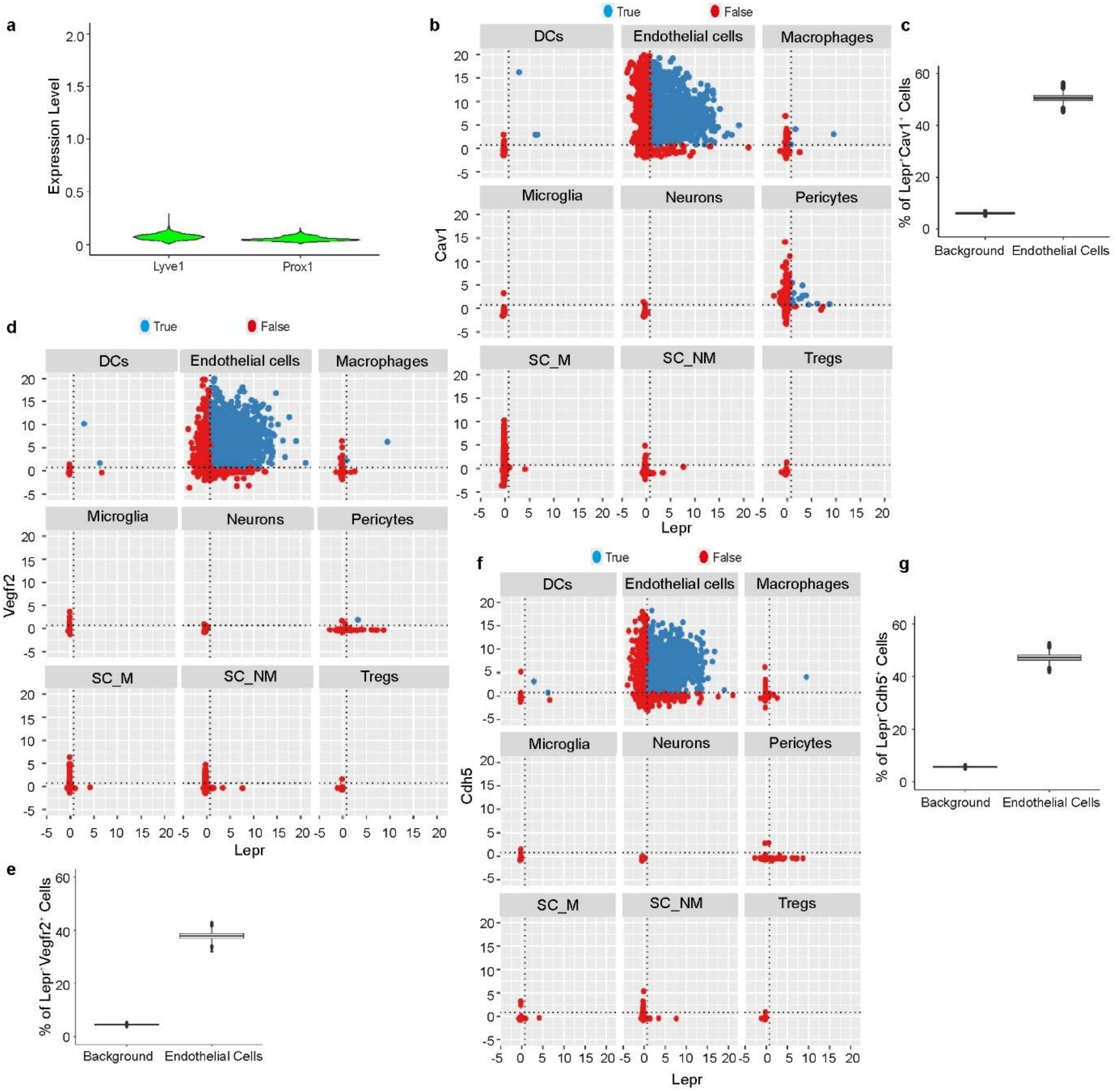


**Extended Data Fig. 2| Identification of Lepr$^+$ vascular endothelial cell population in sympathetic ganglia. a,** Violin plot to show Lyve1 and Prox1 expression in sympathetic endothelial cell population. For visualization, denoised expression data was used. **b,** Correlogram to show the co-expression of Lepr and Cav1 in different cell populations. The position of the dotted lines is described in material and methods. **c,** Box plot shows the percentage of cells co-expressing Lepr and Cav1 in endothelial cluster. **d,** Correlogram to show the co-expression of Lepr and Vegfr2 in different cell populations. The position of the dotted lines is described in material and methods. **e,** Box plot showing the percentage of Lepr$^+$Vegfr2$^+$ endothelial cell population. **f,** Correlogram to show the co-expression of Lepr and Cdh5 in different cell populations. The position of the dotted lines is described in material and methods. **g,** Box plot shows the percentage of cells co-expressing Lepr and Cdh5 in endothelial cluster. Sc_NM = schwan cell_non myelinating, Sc_M = schwan cell_myelinating, DCs = dendritic cells, Tregs = T regulatory cells.

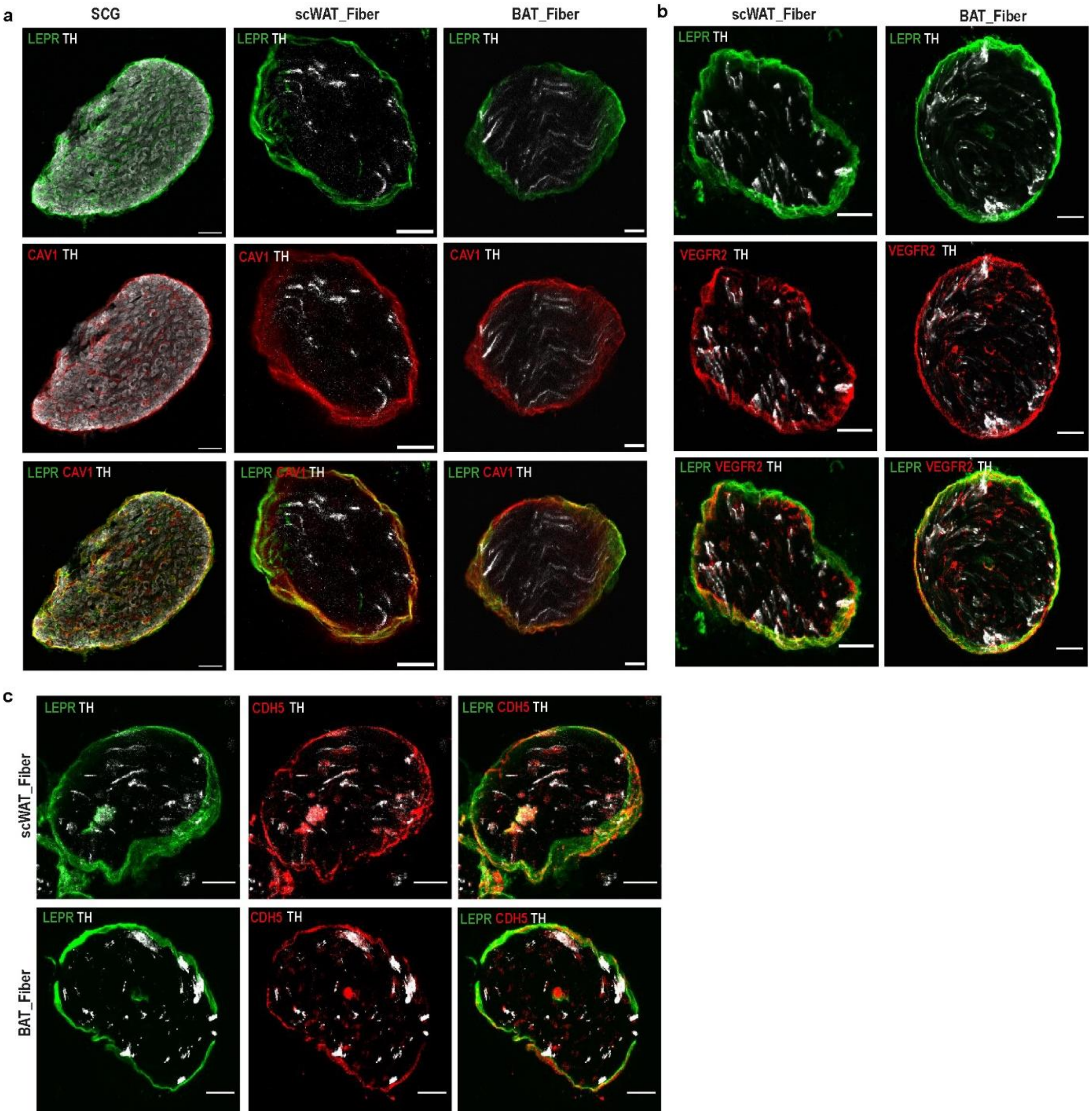


**Extended Data Fig.3|Lepr$^{+}$ sympathetic endothelial cells constitute a thin layer surrounding SCG and sympathetic nerve bundles. a,** Immunofluorescence staining of superior cervical ganglia (SCG) and sympathetic nerve bundles dissected from subcutaneous white adipose tissue (scWAT) and brown adipose tissue (BAT) of LepR$^{cre}$; LSL-YFP reporter adult male mice, showing the expression of LEPR (YFP), CAV1 and TH. **b,** Immunofluorescence staining of sympathetic nerve bundles dissected from scWAT and BAT of LepR$^{cre}$; LSL-YFP reporter adult male mice, showing the expression of LEPR (YFP), VEGFR2 and TH. **c,** Immunofluorescence staining of sympathetic nerve bundles dissected from scWAT and BAT of LepR$^{cre}$; LSL-YFP reporter adult male mice, showing the expression of LEPR (YFP), CDH5 and TH. Scale bar: SCG, 100 µm and Bundles, 20 µm.

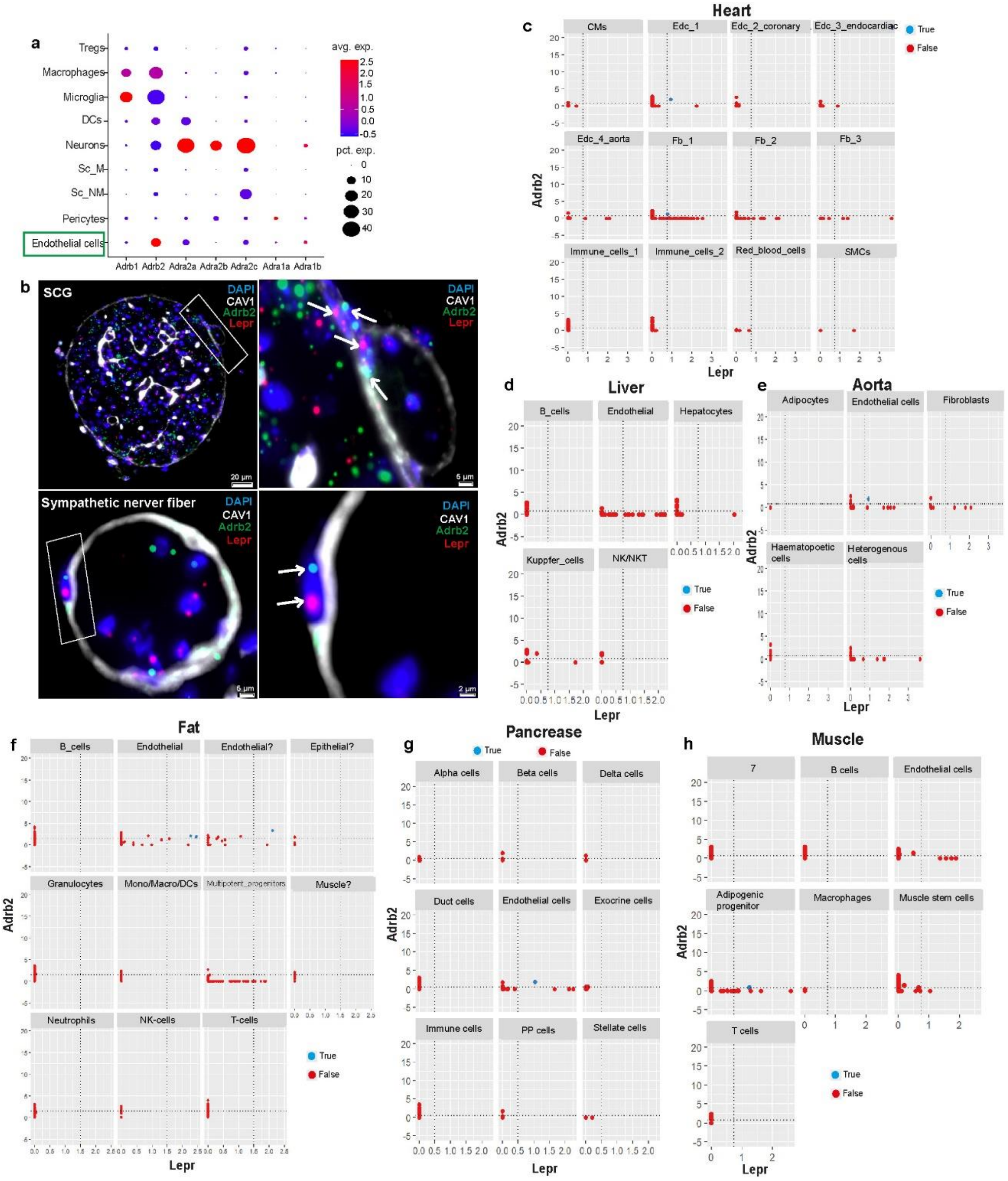


**Extended Data Fig.4|Lepr⁺ SPCs express beta 2 adrenergic (Adrb2) receptor and no other cells in other organs show Lepr and Adrb2 co-expression. a,** Dotplot showing the expression of adrenergic receptors (Adrb1, Adrb2, Adra2a, Adra2b, Adra2c, Adra1a and Adra1b in scRNA sympathetic ganglia dataset. The size of the dot corresponds to the percentage of cells expressing the gene in each cluster and the color represents the average gene expression level. **b,** High magnification image showing the dual labelling of GLUT1 protein with IHC and Lepr, Adrb2 mRNA with ISH in superior cervical ganglia (SCG) and sympathetic nerve bundles dissected from WT adult mice. **c-h,** Correlogram to show the co-expression of Lepr and Adrb2 in different cell population of murine heart (**c**), liver (**d**), aorta (**e**), fat (**f**), pancreas (**g**) and muscle (**h**) extracted from Tabula Muris consortium. The position of the dotted lines is described in material and methods.

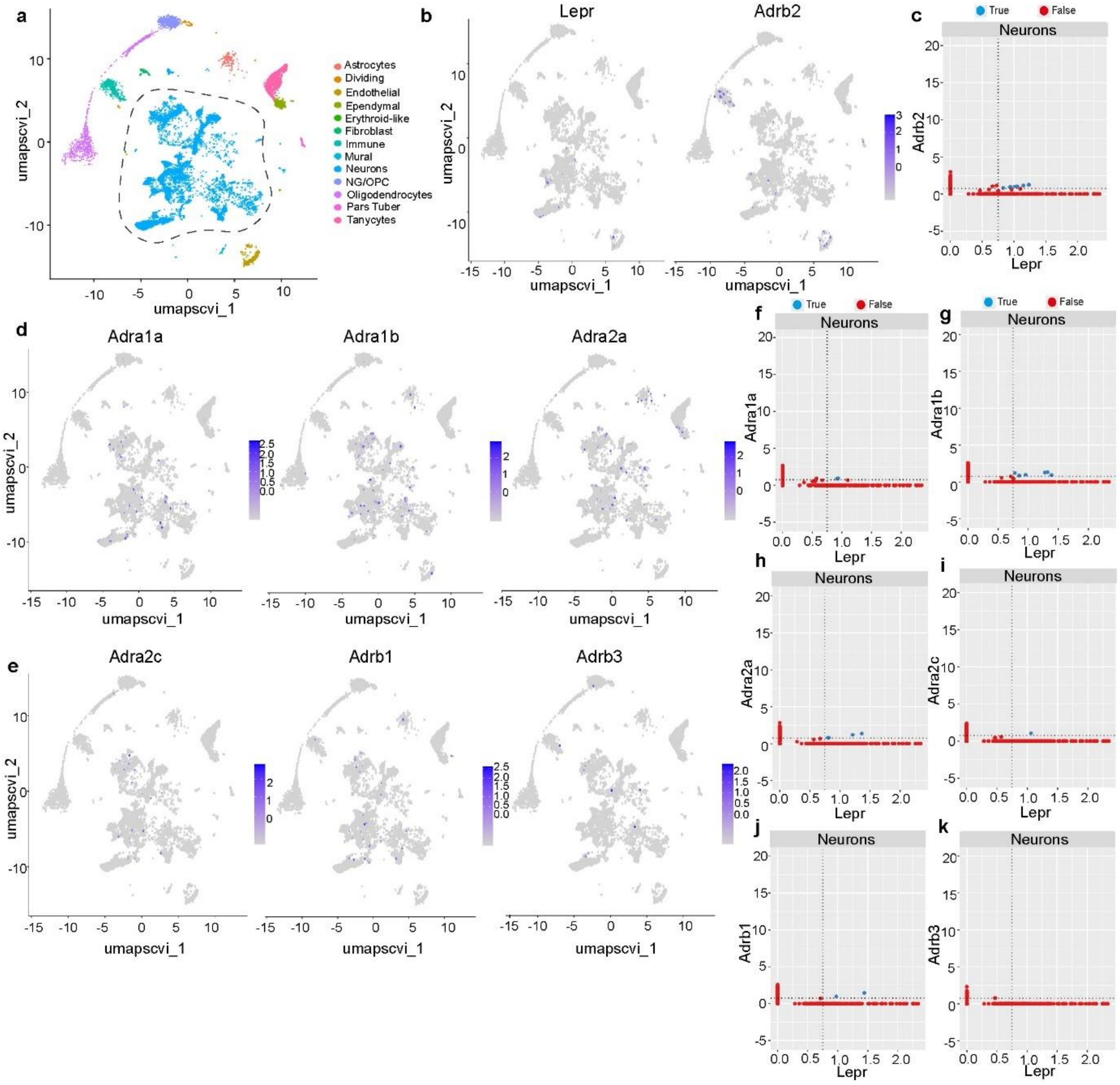

**Extended Data Fig. 5|Leptin receptor and adrenergic receptor co-expression analysis in arcuate hypothalamic scRNA dataset extracted from HypoMap. a,** UMAP plot to show the major cell populations in arcuate hypothalamus. **b,** UMAP plot to show the expression of Lepr and Adrb2 in different cell clusters of arcuate hypothalamus. **c,** Correlogram shows the co-expression of Lepr and Adrb2 in neuronal population of arcuate hypothalamus. **d-e,** UMAP plot to show the expression of other adrenergic receptors (Adra1a, Adra1b, Adra2a, Adra2c, Adrb1 and Adrb3) in different cell clusters of arcuate hypothalamus. **f-k,** Correlogram to show the co-expression of Lepr and other adrenergic receptors in neuronal population of arcuate hypothalamus. The position of the dotted lines is described in material and methods.

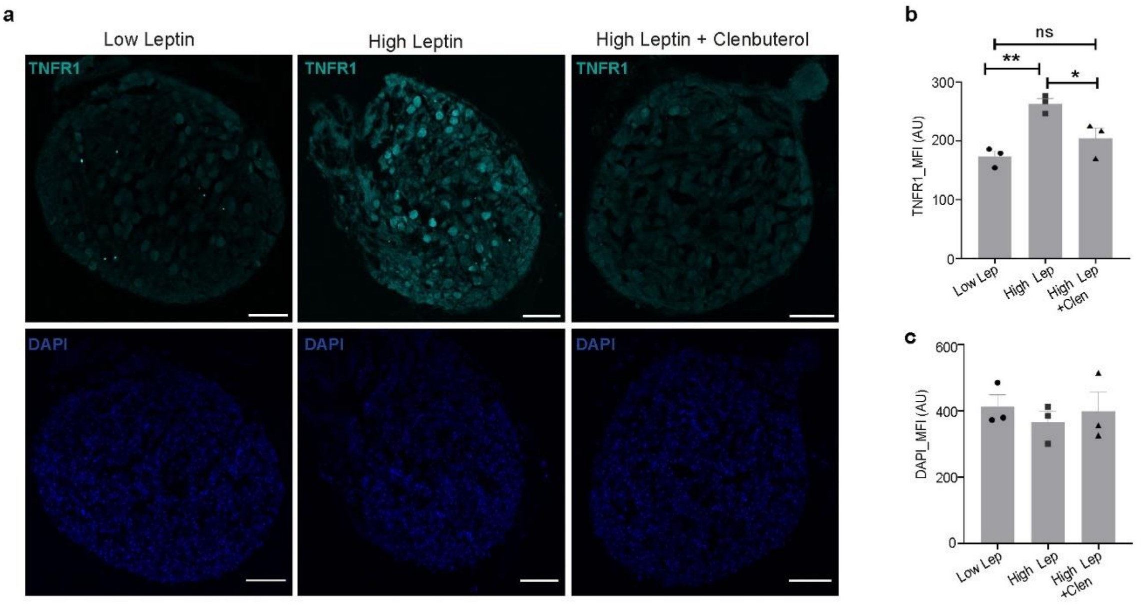


**Extended Data Fig.6|Hyperleptinemia induced apoptosis is reversed by beta 2 adrenergic agonism in SCG explant. a,** Representative images of SCG explant culture showing the expression of TNFR1 and DAPI following low leptin (10 ng/ml), high leptin (100 ng/ml) and high leptin (100 ng/ml) plus clenbuterol (10 µg/ml). Scale bar: 100 µm. **b,** Quantification of the expression of TNFR1 in SCG explant following different stimulating conditions. **c,** Quantification of the expression of DAPI in SCG explant following different stimulating conditions (n = 3 per condition). MFI = Mean Fluorescence Intensity. Data are mean ± s.e.m and were analyzed using one-way ANOVA with Turkey's multiple comparison test. *p < 0.05, **p < 0.01
